# A longitudinal study of *E. coli* lineages and antimicrobial resistance in Ecuadorian children

**DOI:** 10.1101/2020.11.09.375931

**Authors:** Diana Calderón, Paúl A. Cárdenas, Jay Graham, Gabriel Trueba

## Abstract

The gastrointestinal tract (GIT) constitutes a complex and diverse ecosystem. *Escherichia coli* is one of the most frequently studied and characterized species in the gut ecosystem. Nevertheless, there has been little research to determine their diversity and population dynamics in the intestines of children over time. Many intestinal *E. coli* lineages carry antimicrobial resistance and virulence genes, which have implications in disease and public health. In this one-year prospective study, a fresh fecal sample was obtained from 30 children longitudinally for one year (n = 82 fecal samples). From each stool sample, five *Escherichia coli* colonies were randomly selected to characterize their genotype and phenotypic antimicrobial resistance pattern (n = 405 *E. coli* isolates). We found that the most numerically dominant *E. coli* lineages in children’s intestines were transient colonizers, and phenotypic antimicrobial resistance varied significantly over time, however, ST131 a multi-drug resistant pathogen, and 3 additional STs persisted in a child’s intestine for 3 months or more.

**IMPORTANCE:** The length of residency and numeric dominance of antimicrobial-resistant *E. coli* may affect the extent to which an isolate contributes to the dissemination of antimicrobial resistance. We studied the persistence of numerically dominant and antimicrobial-resistant lineages of *E. coli* in the human intestine and found that *E. coli* lineages in the gut of children change rapidly over time.

## INTRODUCTION

*Escherichia coli* is a minor component of the intestinal microbiota of warm-blooded animals (1, 4, and 11). Within the human gut and other warm-blooded animals, there are more than 10 different lineages of *E. coli* at any time; each of these lineages is present in different abundances and remains in the intestine for different periods. Intestinal *E. coli* could be classified as a resident when they remain in the intestine for months and even years (6), transient when they remain for days or weeks (6); dominant when the isolates make up a large proportion (>50%) (26) of the *E. coli* cells in the intestine, and minority when the proportion is smaller (< 10%) (51).

Some *E. coli* are a source of problematic antimicrobial resistance genes (ARGs) (4, 21, and 29) and certain *E. coli* lineages are a major cause of diseases (36, 37, and 38). The abundance of a pathogenic *E. coli* lineage in the intestine has been suggested to be a critical risk factor for intestinal disease (54). Similarly, the abundance of AMR *E. coli* and its persistence in the intestine could be critical in the dissemination of ARGs; *E. coli* is probably the intestinal species that can be transmitted the most between hosts (21, 30 and 34) and it is very active in horizontal gene transfer of ARGs (5, 35 and 37). Here we screened the dominant *E. coli* strains obtained from children less than five years of age and analyzed the turnover of these strains every 3 months for one year. Dominant *E. coli* were defined as the *E. coli* colonies that grew on MacConkey agar at the highest numbers (26).

## RESULTS

We followed over one year a total of 30 children: 8 children with two sampling points (SP I and SP II), and 22 children with three sampling points (SP I, SP II, and SP III). We analyzed 405 *E. coli* isolates from 82 fecal samples from 30 individuals. The large majority of fecal samples (81.71%) contained one or 2 dominant *E. coli* isolates (isolates carrying 1 or 2 different *fumC* alleles) (Table S1 and Table S2). When we analyzed the diversity of all the isolates, we found that 64% of fecal samples contained isolates with the same *fumC* alleles: 34% of fecal samples had isolates with the *fumC11* allele, 11 (13%) fecal samples had isolates with *fumC35* and 9 (11%) fecal samples had isolates with *fumC4* allele. Among the 405 isolates, we found a total of 41 *fumC* alleles; the most prevalent was *fumC11* (n=108, 26.7%), followed by *fumC35* (n=40, 9.8%) and *fumC4* (n=38, 9.4%). We found only 4 individuals who carried isolates of the same ST in fecal samples: ST34 was the predominant genotype recovered from one child (Child #26, isolates recovered 11 months apart), ST10 was the predominant genotype in two children (Child #20, isolates recovered 7 months apart) and (Child # 30, isolates recovered 5 months apart), and ST131 was the predominant genotype in one child (Child #23, isolates recovered 4 months apart) (Fig. 1).

**FIG 1.**
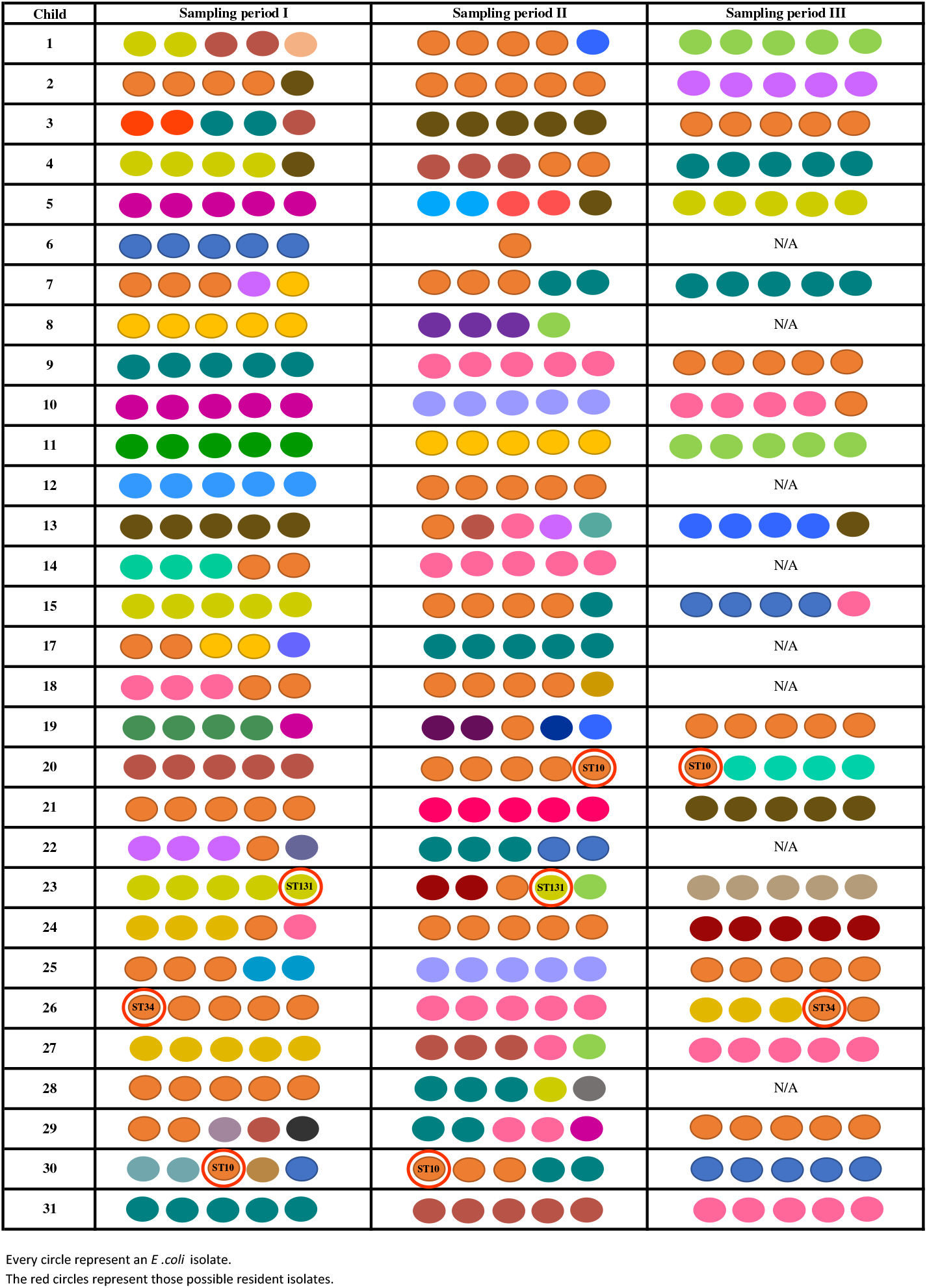
Diversity of numerically dominant fecal *E. coli* isolates obtained during 3 sampling periods Different colors indicate different *fumC* alleles and ST numbers indicate strains belonging to sequence types found in different sampling periods

Among the 405 isolates recovered, 122 (30.1%) were susceptible to all the antimicrobials tested. The remaining fell into one of the 76 unique antibiotic resistance profiles: 50 isolates (12.3%) were resistant to only one antibiotic, 37 isolates (9.1%) were resistant to two antibiotics, and 196 (48.4%) were resistant to three or more antimicrobials.

Phenotypic resistance to the combination of Ampicillin (AM), Trimethoprim-sulfamethoxazole (SXT), and Tetracycline (TE) was the most common profile (n=36, 8.9%), followed by Tetracycline (TE) resistance (n=21, 5.2%) and Ampicillin (AM), Trimethoprim-sulfamethoxazole (SXT), and Amoxicillin-clavulanic acid (AMC) resistance (n=15, 3.7%).

Among the eight children who were enrolled for SP I and SP II, the highest percentages of resistance in SP I isolates was to Tetracycline (TE) (50%), Trimethoprim-sulfamethoxazole (SXT) (37.5%), and Ampicillin (AM) (35%), while in SP II isolates was to Ampicillin (AM) (54.3%), Trimethoprim-sulfamethoxazole (SXT) (40%) and Tetracycline (TE) (31.4%). There were significant statistical differences in the resistance to Chloramphenicol (C) between SP I and SP II, with resistance higher in SP I (Table 1 and Table S3).

**TABLE 1.** Antimicrobial susceptibility profiles of *E. coli* isolates obtained from children (fecal samples) in different sampling periods (SPI, SPII, SPIII).

| Antimicrobials | n= 110 isolates (each one) |  |  | Chi square test | P value |
| --- | --- | --- | --- | --- | --- |
|  | SP I | SP II | SP III |  |  |
| Ampicillin (AM) | 64 (58.2 %) | 60 (54.5 %) | 46 (41.8 %) | **6.502** | **0.039** |
| Chloramphenicol (C) | 5 (4.5 %) | 12 (10.9 %) | 9 (8.2 %) | 3.090 | 0.213 |
| Imipenem (IPM) | 12 (10.9 %) | 1 (0.9 %) | 7 (6.4 %) | **9.687** | **0.008** |
| Trim-sulfamet (SXT) | 58 (52.7 %) | 58 (52.7 %) | 50 (45.4 %) | 1.552 | 0.460 |
| Gentamicin (GM) | 0 (0 %) | 6 (5.4 %) | 0 (0 %) |  |  |
| Ceftazidime (CAZ) | 2 (1.8 %) | 14 (12.7 %) | 10 (9.1 %) | **9.352** | **0.009** |
| Cefepime (FEP) | 13 (11.8 %) | 15 (13.6 %) | 9 (8.2 %) | 1.705 | 0.426 |
| Cefazolin (CZ) | 18 (16.4 %) | 14 (12.7 %) | 15 (13.6 %) | 0.645 | 0.724 |
| Ciprofloxacin (CIP) | 22 (20 %) | 21 (19.1 %) | 9 (8.2 %) | **7.168** | **0.028** |
| Cefotaxime (CTX) | 14 (12.7 %) | 18 (16.4 %) | 14 (12.7 %) | 0.808 | 0.668 |
| Amoxicillin + Clav. Ac.(AMC) | 35 (31.8 %) | 28 (25.4 %) | 15 (13.6 %) | **10.376** | **0.006** |
| Tetracycline (TE) | 55 (50 %) | 52 (47.3 %) | 50 (45.4 %) | 0.462 | 0.794 |

(P value < 0.05)

Among SP I – SP III, which included 22 children, the highest percentages of resistance in SP I and SP II isolates corresponded to Ampicillin (AM) (SP I=58.2% and SP II=54.5%), Trimethoprim-sulfamethoxazole (SXT) (SP I and SP II=52.7%) and Tetracycline (TE) (SP I=50% and SP II=47.3%), while in SP III isolates were to Trimethoprim-sulfamethoxazole (SXT) (45.4%), Tetracycline (TE) (45.4%) and Ampicillin (41.8%) (Table 1 and Table S3).

## DISCUSSION

During the current prospective study, we followed 30 children for 2 to 3 sampling periods (for 1 year) and screened the numerically dominant *E. coli* strains present in fresh fecal samples. At least three of the five selected colonies from each sample showed the same *fumC* allele and the same antimicrobial resistance profile, providing additional evidence that the isolates were numerically dominant in the individual (Figure 1). We observed that very few strains (n=4) persisted over time in some individuals, suggesting that most numerically dominant strains are transient colonizers and that there is a high turnover rate of *E. coli* lineages. This observation is in agreement with previous reports showing a high diversity and high turnover rate of *E. coli* lineages in the human intestines [10, 18, 53]. The high turnover rate may be due to bacteriophage, bactericins, protozoal predation, or immune mechanisms [4, 10].

Recent evidence has indicated that diet can also favor the proliferation of some strains of other intestinal bacteria [35]. We identified only four children with possible dominant resident strains (i.e. strains that persisted throughout the study period): ST10 (individuals 20 and 30), ST34 (individual 26), and ST131(individual 23), isolates which persisted for 7, 5, 11, and 4 months, respectively. The most remarkable of them is ST131 which is a known human extra-intestinal pathogen, characterized by its virulence (37) and resistance to extended-spectrum cephalosporins (ESC) and fluoroquinolones (FQ) (37 and 39). This ST is strongly related to extra-intestinal infections, mainly urinary tract (10 and 13) which has been reported as dominant in human intestines (53), domestic animals intestine the environment (36–39). Moreover, ST10 and ST34 belonged to clonal complex 10 (CC10); ST10 is known as one of the most dominant commensal STs in *Escherichia coli* (38), however, strains belonging to this ST are often not clonal (40).

According to the surveys, 30 households reported access to potable water 24/7, and 28 had sewerage. During the sampling periods, 20 children changed their dietary habits (stop breastfeeding), 14 children were exposed to antimicrobials at least in one sampling period, and 29 had contact with domestic animals (pets, livestock, poultry, etc.), and 14 children had diarrhea at least in one sampling period. These events may have altered the frequency of some of the *E. coli* lineages, however, it is difficult to assess their impact (45).

Regarding phenotypic antimicrobial resistance, among the 405 isolates analyzed, 30.1% (122 of 405) were susceptible to all of the antimicrobials tested. Resistance to Ampicillin (AM), Trimethoprim-sulfamethoxazole (SXT), and Tetracycline (TE) were the most prevalent, while Gentamycin (GM) was the least because only six strains showed resistance. We found several commensal *Escherichia coli* isolates resistant to Imipenem (IPM) (24 out of 405, 5.9%) and Cefepime (FEP) (40 out of 405, 9.8%) which are resistances that should be associated with hospitals and not community isolates (41).

There were significant statistical differences in resistance to Ampicillin (AM) between SP I and SP II (p=0.004, Table 1), and resistance to Ampicillin (AM), Ceftazidime (CAZ), Ciprofloxacin (CIP), and Amoxicillin + clavulanic acid (AMC) was significantly different between SP I, SP II and SP III (p= 0.039, 0.009, 0.028, and 0.06, respectively) (Table 2). Our findings suggest that prevalence surveys of AMR in *E. coli* are potentially not that useful because the phenotypic resistance profiles change rapidly over time, without any strong driving factors. Understanding the factors involved in lineage turnover is critically important for understanding antimicrobial resistance and virulence gene carriage in commensal strains.

Our research shows another level of complexity on the understanding of the gut microbiome, which is important as many strains of the same species carry different metabolic, virulence, or antibiotic resistance genes. Our study reinforces the idea of the constant transition of microbiome members over time [4, 5, 11, 12, and 19] and the need for more research in the dynamics of strain populations in the intestine.

## MATERIALS AND METHODS

### Study locations

This one-year longitudinal study was carried out in six semi-rural communities belonging to the parishes of Yaruquí, Pifo, Tumbaco, Checa, Puembo, and Tababela; all of them located near Quito, Ecuador. For household enrollment, inclusion criteria included: (i) households have a child ages six months to four years, (ii) households have a childcare provider who was over eighteen years, and (iii) informed consent was provided by the primary childcare provider. Sixty-one households were enrolled at the beginning of the study but, only 27 finished the full longitudinal study.

### Ethical considerations

The study protocol was approved by the Committee for Protection of Human Subjects (CPHS) and the Office for Protection of Human Subjects (OPHS) at the University of California, Berkeley (Federalwide Assurance # 6252), the Human Research Ethics Committee at the Universidad San Francisco de Quito (no. 2017-178M) and the Ministry of Public Health, Ecuador (MSPCURI000243-3).

### Sample collection

We collected a single fecal sample from each child during three sampling periods (SP): from October to December of 2018 (SP I), from January to May of 2019 (SP II), and from July to December of 2019 (SP III), obtaining a total of 120 stool samples. Each time a sample was collected, the childcare provider completed a survey related to the current family lifestyle and recent exposure-related factors relevant to AMR. Eighty-one stool samples were obtained from the same 27 children along the three sampling periods. Moreover, a total of 10 fecal samples were obtained from the same five children in the sampling periods I and II. The remaining 29 stool samples were obtained only during sampling period I; therefore, those samples were discarded for further analyses. Each stool sample was collected into sterile tubes, stored at 4°C, transported to the laboratory, and immediately processed.

### *Escherichia coli* isolation

Each fecal sample was plated on MacConkey agar and incubated at 37 °C for 18 hours. To ensure the selection of the dominant *Escherichia coli* strain (≥ 50% of all colonies) [26], we collected 5 colonies that correlated with the predominant colony morphology. Additionally, each colony was transferred to Chromocult® coliform agar for the identification of *Escherichia coli* through its β-D-glucuronidase activity. Those strains were incubated in Brain Heart Infusion (BHI) medium + glycerol (15%) at 37 °C for 18 hours to perform the antimicrobial susceptibility test. After that, the tubes were stored at −80 °C.

### Antimicrobial susceptibility test

We used the Kirby Bauer technique (disc diffusion in Muller Hinton agar) to determine the strains antimicrobial susceptibility using the following twelve antimicrobial discs: Cefazolin (CZ; 30 μg), Ciprofloxacin (CIP; 5 μg), Ampicillin (AM; 10 μg), Chloramphenicol (C; 30 μg), Imipenem (IPM; 10 μg), Trimethoprim-sulfamethoxazole (SXT; 1.25/23.75 μg), Gentamicin (GM; 10 μg), Ceftazidime (CAZ; 30 μg), Cefepime (FEP; 30 μg), Cefotaxime (CTX; 30 μg), Tetracycline (TE; 30 μg) and Amoxicillin + Clavulanic Acid (AMC; 20/10 μg). Resistance or susceptibility was determined according to Clinical and Laboratory Standards Institute (CLSI) guidelines [41].

### DNA extraction

Each isolate was grown on MacConkey agar at 37 °C for 18 hours and 5-6 colonies were placed into Eppendorf® tubes with 500 μl of sterile distilled water and DNA was released by boiling cells suspensions for 1 minute. The quality of the DNA was monitored by gel electrophoresis.

### Strains Genotyping

The clonal relationship of the isolates was screened by amplifying and sequencing the *fumC* gene [21] using the Master Mix Go Taq. Those isolates coming from the same individual and sharing the same allele were subjected to full multi-locus sequence-typing (MLST). Briefly, PCR conditions were: 180 secs at 95°C, 30 cycles of 30 secs at 94°C, 30 secs at the annealing temperature of each primer *(adk:* 52 °C; *fumC:* 55 °C; *gyrB* and *mdh:* 58 °C; *icd* and *recA:* 54 °C;*purA:* 50 °C) and 60 secs at 72 °C *(fumC* and *icd)* or 45 secs at 72 °C *(adk, gyrB, mdh, purA,* and *recA*), and a final extension of 7 minutes at 72 °C.

### DNA sequencing

All PCR products were sequenced at Macrogen Inc. using the Sanger sequencing method. The sequences were analyzed using the program Geneious Prime 2020 and were screened in the Enterobase database [50].

### Statistical analysis

Significant differences between phenotypic antimicrobial resistance prevalence of the individuals through time were tested using a chi-square test.

## ACKNOWLEDGMENTS

P.C. is funded by NIH FIC D43TW010540 Global Health Equity Scholars. The study was also supported by the National Institutes of Health under Award Number R01AI135118. The content is solely the responsibility of the authors and does not necessarily represent the official views of the National Institutes of Health. The funders had no role in study design, data collection, and interpretation, or the decision to submit the work for publication.

